# Neural activity precedes conscious awareness of being in or out of a transient hallucinatory state

**DOI:** 10.1101/2020.03.17.995282

**Authors:** Kenneth Hugdahl, Alexander R. Craven, Erik Johnsen, Lars Ersland, Drozdstoy Stoyanov, Sevdalina Kandilarova, Lydia Brunvoll Sandøy, Rune A. Kroken, Else-Marie Løberg, Iris E. Sommer

## Abstract

Auditory verbal hallucinations, or “hearing voices”, is a remarkable state of the mind, occurring in psychiatric and neurological patients, and in a significant minority of the general population. An unexplained characteristic of this phenomenon is that it transiently fluctuates, with coming and going of episodes with time. We monitored neural activity with BOLD-fMRI second-by-second before and after participants indicated the start and end of a transient hallucinatory episode during the scanning session by pressing a response-button. We show that a region in the ventro-medial frontal cortex is activated in advance of conscious awareness of going in or out of a transient hallucinatory state. There was an increase in activity initiated a few seconds before the button-press for onsets, and a corresponding decrease in activity initiated a few seconds before the button-press for offsets. We identified the time between onset and offset button-presses, extracted the corresponding BOLD time-courses from nominated regions-of-interest, and analyzed changes in the signal from 10 seconds before to 15 seconds after the response-button was pressed, which identified onset and offset events. We suggest that this brain region act as a switch to turn on and off a hallucinatory episode. The results may have implications for new interventions for intractable hallucinations.

## Introduction

Auditory verbal hallucinations (AVH) in the sense of “hearing voices” in the absence of a corresponding auditory source, is a remarkable state of the mind. AVHs were traditionally seen as a hallmark of schizophrenia^1–7^, but also occur in other psychiatric and neurological disorders. AVH cross the border between pathological and normal states of mind, since they are experienced in about 10% of the general population^8–11^. As a symptom, AVHs are often experienced as highly distressing, while people in the general population are usually not distressed to the same degree^12, 9^. An equally remarkable characteristic of AVHs is that they are spontaneous episodes for which there are no known environmental triggers, occurring in resting and relaxed as well as in stressful and noisy environments^13, 14^. The same is true for the offset of an episode, which likewise can occur in a variety of environmental situations. The absence of environmental causes for these transient on- and off-fluctuations of AVHs would thus point to an internal, i.e. some kind of neural switching mechanism, not only for the spontaneous onset, but also for the spontaneous offset of an episode. We therefore studied the neural underpinnings of the spontaneous switching between AVH on- and off-states by monitoring neural activity a few seconds before and after reported onset (start) and offset (stop) of a hallucinatory episode, and related this to the corresponding conscious awareness of the event. Brain activity can be monitored on-line with functional magnetic resonance imaging (fMRI), where changes in neural activity are estimated from modelling of the bloodoxygenation-dependent (BOLD) function^15^. Yet this requires participants to lay still in the scanner while experiencing AVH and indicating their on and offset by button-press, a very demanding task (see ^9,16–19^). This paradigm requires patients to be aware and thoughtful of their experiences and to experience a required minimum of episodes of AVH during the scanning session, as neither continuous hallucinations nor a period with too few hallucinations will allow the study of on- and offsets. Only few research groups around the world have succeeded to obtain a number of such scans, typically less than 20. In order to increase power to detect subtle changes in activity during brief moments of on and offsets, we joined forces from three research groups to recruit a reasonably large sample of individuals who were hallucinating frequently, but not continuously. The aim of the present study was to use fMRI to monitor changes in neural activity on a second-by-second-basis, in a resting-state situation, where subjects signaled the onset of a hallucinatory episode by pressing one response-button and the offset of an episode by pressing another response-button, in the course of the scanning session. A time-window was set from 10 seconds before to 15 seconds after the subject had pressed a button, from which voxel-wise data were extracted, analyzed and displayed second-by-second in a sliding window over the evaluation period (see Methods). Data were pooled from three different sites at the University of Bergen, Norway, Groningen University Medical Center, Netherlands, and Medical University of Plovdiv, Bulgaria, making up a total of 66 subjects hallucinating during fMRI recordings. As hallucinatory experiences cross the border between abnormal and normal conditions, we included both clinical and non-clinical “voice-hearers”, focusing on tracking the neural signatures of AVH-experiences per se, not restricted to a particular diagnostic group or mental condition.

## Results

### Anticipated activity patterns during hallucinatory periods

Functional data that were obtained using the button-press symptom-capture paradigm^9^ revealed several statistically significant clusters of increased activity during hallucinatory periods (i.e. periods after onset and before offset). These clusters included the left fronto temporal language areas (Wernicke’s area in the superior temporal gyrus, and Broca’s area in the inferior frontal gyrus), using a FEW-corrected significance threshold of p <.05 (see Figure 1 and Table 1).

**Figure 1.**
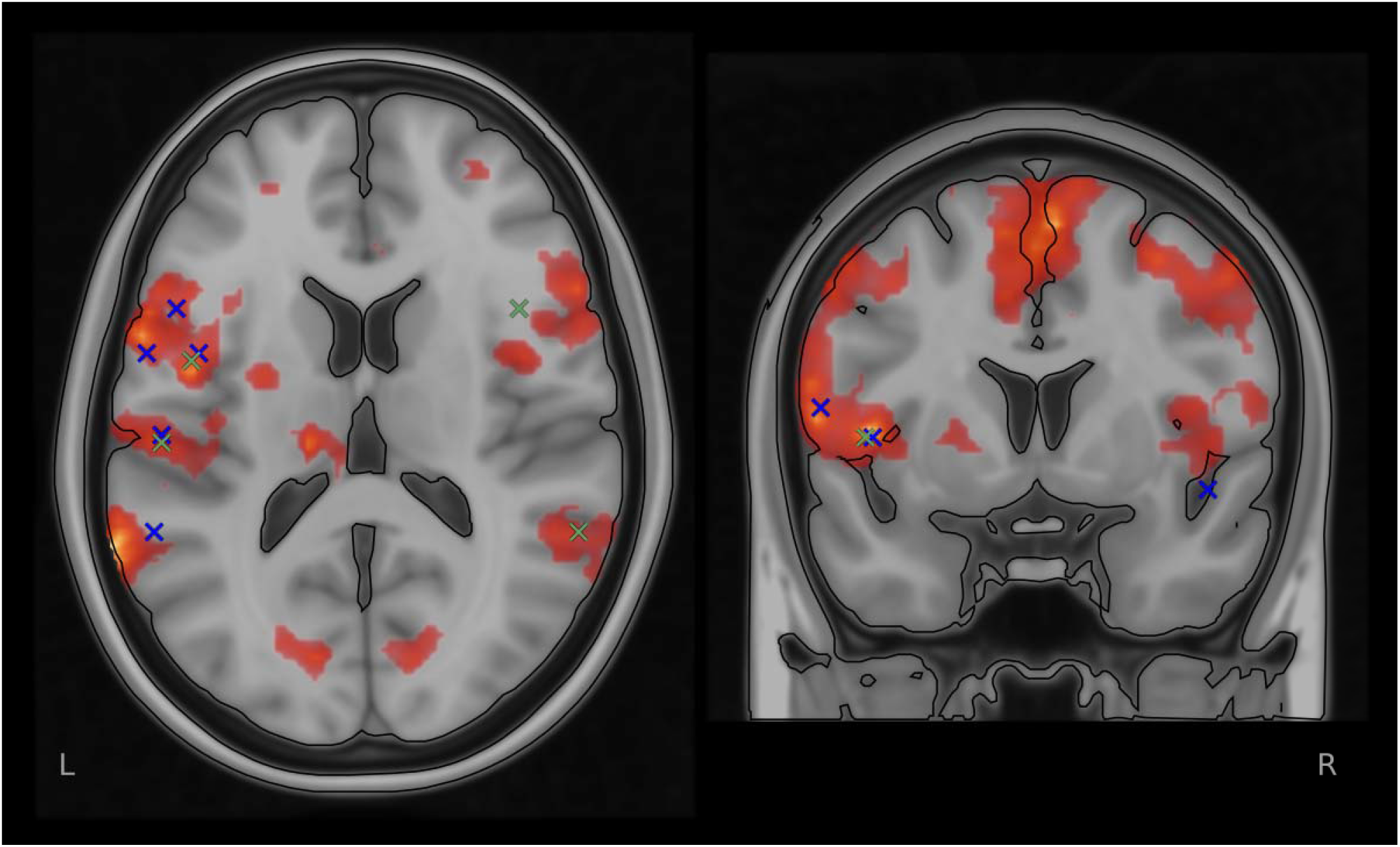
shows the results from the standard group-level block-analysis of the BOLD-fMRI data, overlaid with peak activity from the Jardri, et al.^16^ and Kompus, et al.^17^, meta-analyses, marked respectively with a blue (Jardri) and green (Kompus) ‘x’, verifying the presently seen activity with activity previously repeatedly reported in the literature.

**Table 1.**
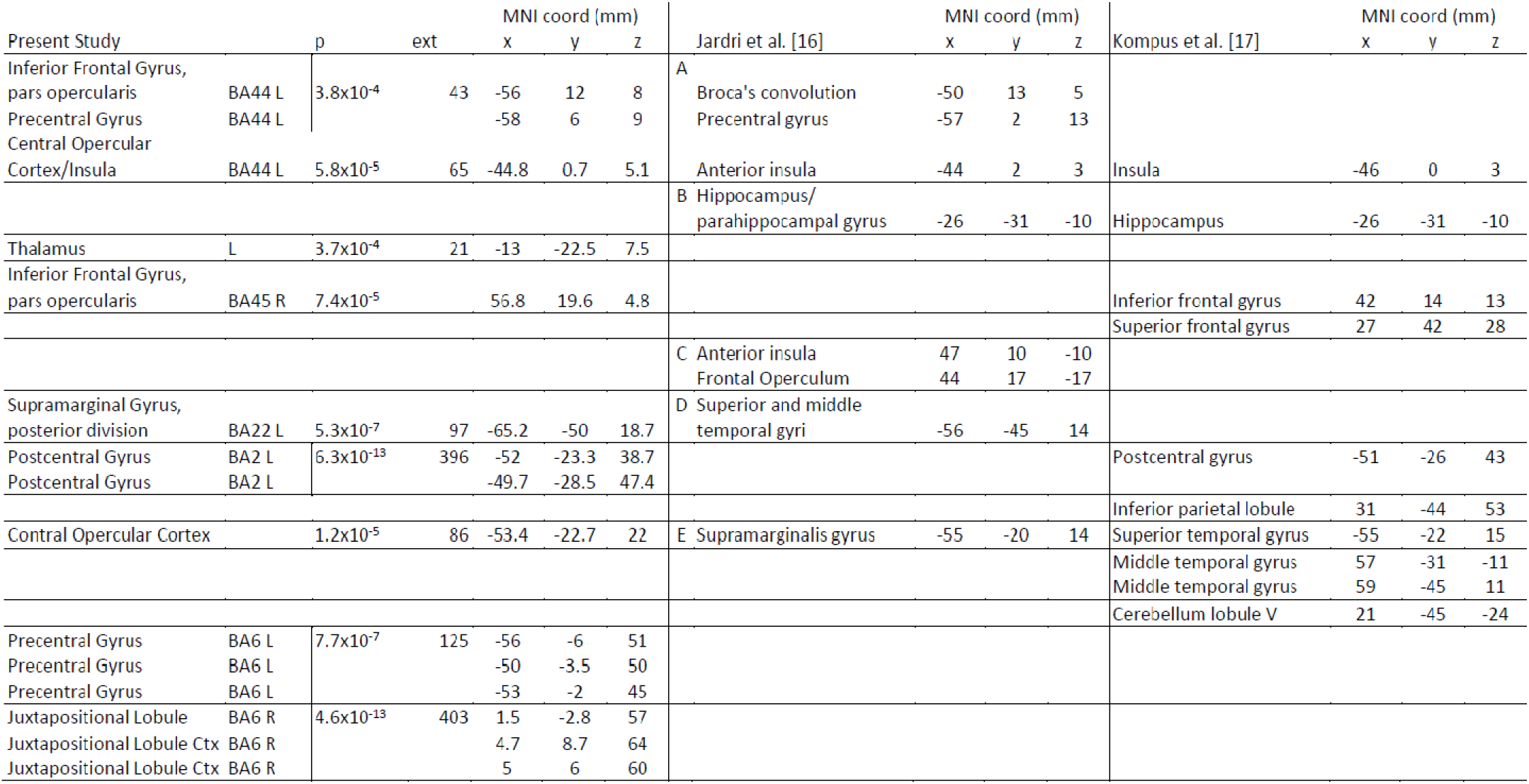
shows clusters and local maxima obtained by block analysis in the present study, with corresponding activations from Jardri, et al.^16^ and Kompus, et al.^17^. Clusters in the novel data are thresholded non-parametrically at Z>4.5; corrected cluster significance is reported, thresholded at p>0.05 and extent > 20 voxels.

We compared the anatomical localizations of these activity with areas previously reported as activated during ongoing hallucinatory episodes (see meta-analysis^16, 17^). As can be seen in Figure 1, the overlap of the present activity with previous reports is almost perfect, thus validating that the present activity reflects neural correlates of ongoing hallucinations. Analyzing activity data from each of the three sites separately, the general pattern of activity remained, although statistically weaker for the Bergen cohort and reduced to trend level for the Plovdiv cohort. Contrasting clinical- and non-clinical voice hearers from the Groningen cohort revealed overlapping activity in all areas except for the right planum temporale (PT) and right lateral superior occipital cortex.

### Differential activity in the ventro-medial frontal cortex

Analysis of the extracted time-courses from the nominated regions of interest (ROIs), separately for onset and offset of episodes, revealed a distinct decrease in activity in the intersection of the paracingulate cortex, medial frontal cortex, and the frontal pole (see Figure 2). The decrease had a minimum peak at time (t) = 3 sec (Δ = −158 iu, p = 0.021 ±~0.002, 95% CI) relative to the button-press response (see Figure 3). This minimum preceded the corresponding motor response from the subjects’ button-press which had a peak maximum at t = 5 seconds.

**Figure 2.**
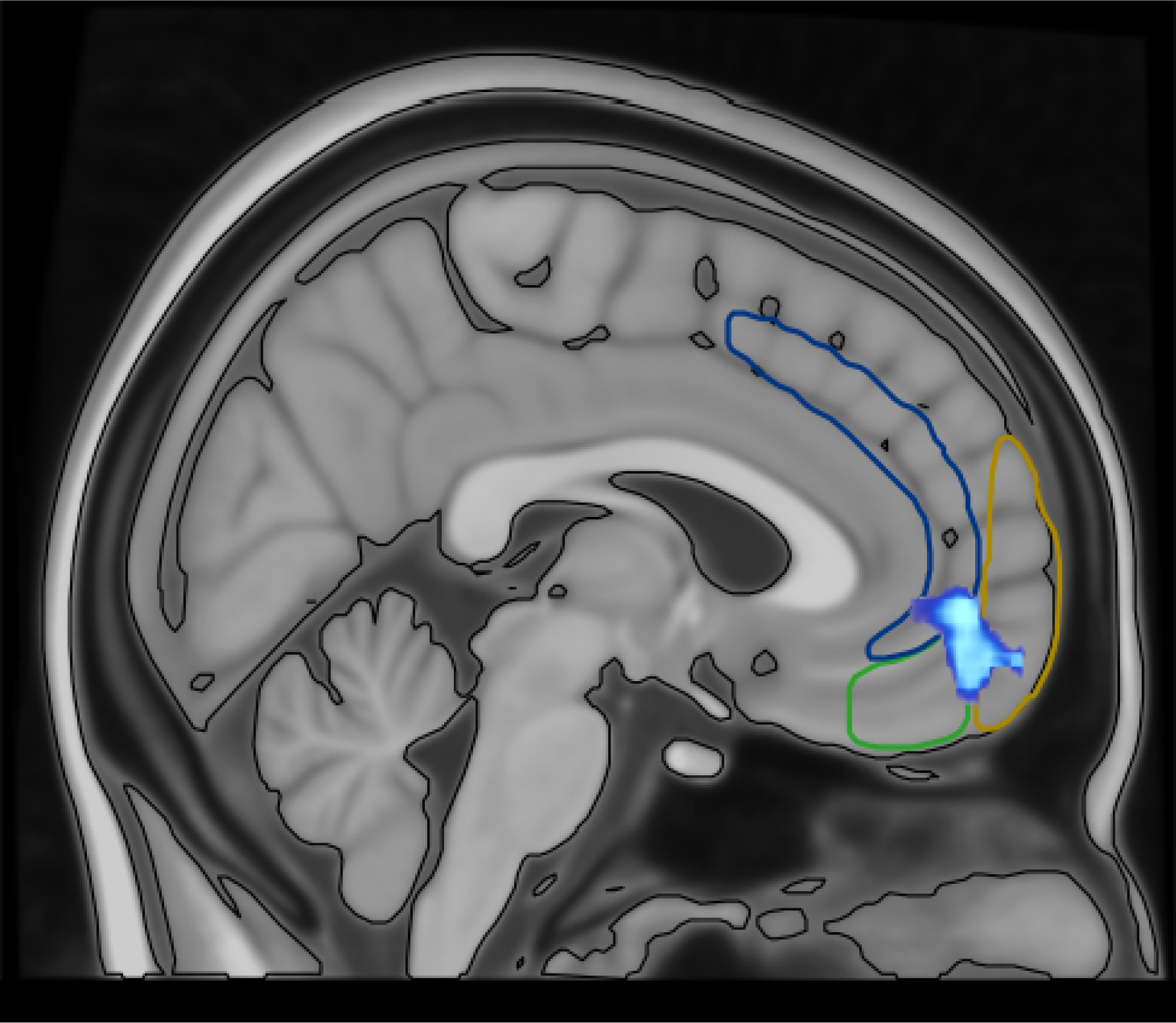
shows the anatomical localization of the ROI (turquoise) with MNI peak coordinates x=8.8, y=44.8, z=-5.64 mm from where the time-courses were extracted, in the intersection of the paracingulate cortex (demarcated in dark blue), medial inferior frontal cortex (demarcated in green) and frontal pole (demarcated in yellow).

The decrease in activity preceding an offset button-press contrasted to an increase in activity observed in advance of an onset button-press (Δ = 35 iu, p = 0.014, ±~0.0016, 95% CI), occurring at around t = −1 second. These results were verified in an extended time-series analysis (see Figure 4) averaged across subjects, and where aberrant time-courses were rejected and results iteratively updated. An HRF-model fit was thereafter applied to the data, revealing a similar differential pattern for onset- and offset-events as seen in Figure 3. See Methods section for further details.

**Figure 3.**
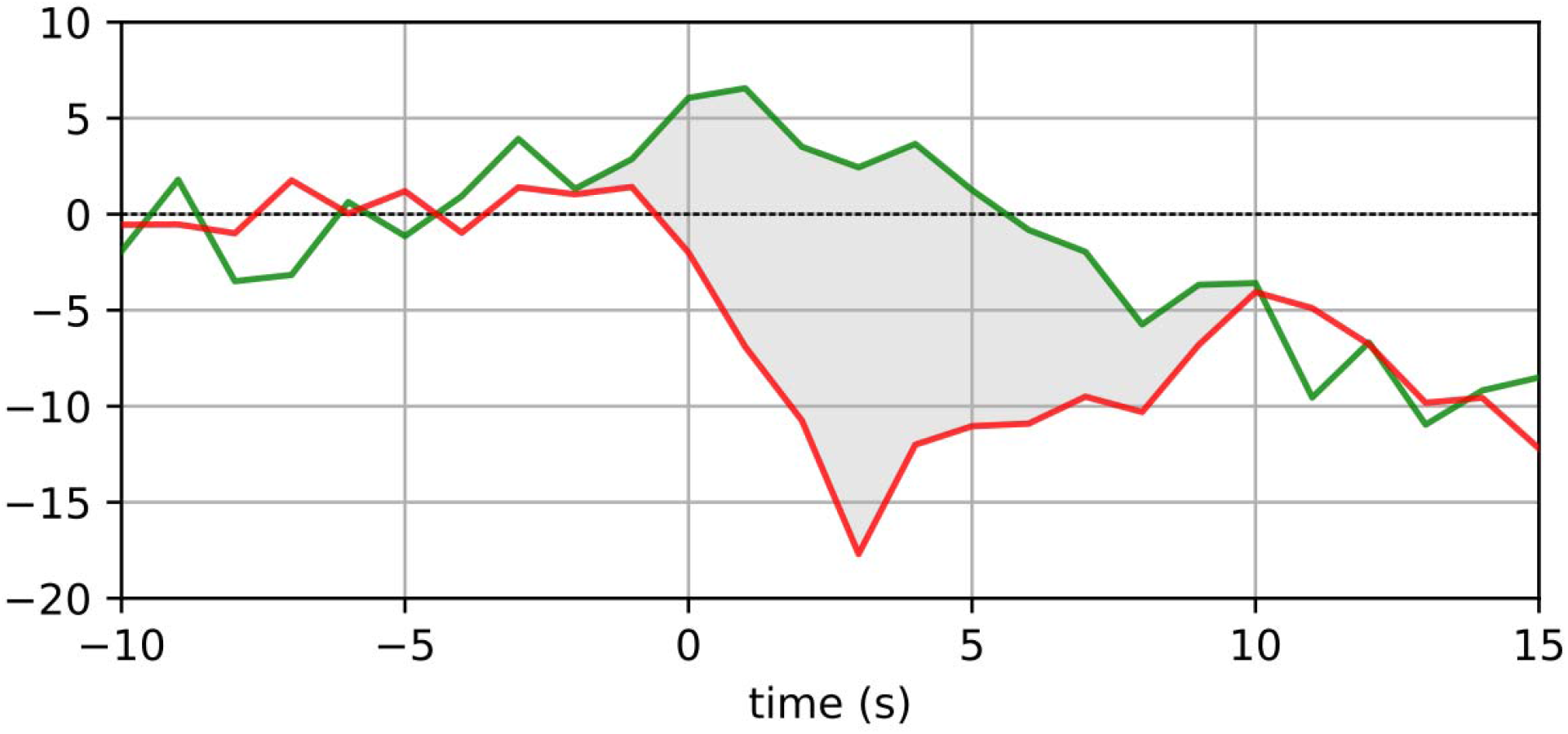
shows filtered time-courses tracked second-by-second around the onset-of-hallucination (red) versus offset-of-hallucination (green) events from the ROI at the intersection of the paracingulate cortex/medial inferior frontal cortex/frontal pole. Grey area shows differential responding for onset-versus offset-events. Time “0” on the x-axis represent the point in time when the button-press occurred. Time −10 represent 10 secs before a button press and time 15 represents 15 sec after a button-press. Y-axis in international units (iu). See Results section for further details.

These results were obtained using the strictest criteria for interpretation of the subjects’ button-press response data, and were maintained (with varying levels of significance) for more liberal interpretations (Δ = −79 iu, p = 0.16, ±~.005, 95% CI, Δ = 36 iu, p = 0.0024, ±~.0007, 95% CI for the long-block interpretation, see Methods for explanation). ROIs in the in the pre-central motor-area also exhibited significantly increased activity, with peak activity at t = 5 seconds; all significances after FEW-correction and p-level set to <.05.

### 4D permutation analysis

Full-volume permutation analysis across the nominated time-windows further confirmed a differential direction of activity for offsets versus onsets at the ventral edge of the intersection of the paracingulate cortex, medial frontal cortex and the frontal pole (see Figure 2). This region exhibited significantly reduced activity (p = 2.5×10^-8^) for offset-of-hallucination events relative to onset-of-hallucination events, peaking 2 seconds after the recorded button-press event, and 3 seconds before the peak of the motor activity associated with the button-press itself. Additional contrasting activity, not directly attributable to the motor response, was observed in the inferior frontal gyrus (p = 1.4×10^-11^, with peak at time (t) = 17 sec), and the central opercular cortex bordering on Heschl’s gyrus (p = 4.6×10^-14^ with peak at time (t) = 19 sec). These regions were also identified in our functional block-analysis (see Table 1), and have also been mentioned in the literature^16, 17^ (see Table 1).

The subsequent permutation analysis of offset-of-hallucination events against “baseline” data from random time-windows showed again significantly reduced activity (p = 5.2×10^-7^) specific to offset-of-hallucinatory events. This analysis also revealed transient increases in activity in the anterior cingulate (p = 3.7×10^-6^), insula/left operculum (p = 9.3×10^-6^), thalamus (p = 1.8×10^-7^) and paracingulate cortex (p = 1.8 × 10^-7^). Permutation analysis with onset-of-hallucinatory events against “baseline” data also revealed a slight increase in activation initiated before the button-press with a peak around 1 second after the button-press event (p = 2.5 × 10^-4^). Finally, there was a large and significant increase in activity in the primary motor cortex in the pre-central gyrus (p = 7×10^-32^), representing the button-press response per se.

## Discussion

The finding of anticipatory neural activity in the ventro-medial frontal region preceding start and end of auditory hallucinations is a novel finding, which could lead to new hypotheses about excitation and in particular inhibition of a hallucinatory event, and what underlies the experience of the start and stop of a “voice”. Both the time-course- and permutation-analyses revealed a significant brain response initiated a few seconds in advance of the subject becoming consciously aware of being in or out of a hallucinatory state. This suggests that the ventro-medial frontal region may be crucial in both the initiation and cessation of hallucinatory episodes, and speaks to a kind of regulatory role, or switch function for this region. Metaphorically, it could be thought of as this region acting like a conductor, directing the orchestration of neural events that leads up to a full-blown perceptual experience of “hearing a voice” in the absence of a corresponding external source and also to the cessation of the perceptual experience. Brain responses in advance of conscious awareness have been previously reported for error monitoring, where subjects showed reduced activity in regions related to the default mode network (the “oops region”) seconds before they became aware of having made an erroneous response^20^. The present results complement recent findings of aberrant functional connectivity^21^ and morphological differences^22^ in the same brain region. Garrison, et al. used structural MRI and found that hallucinating patients had shorter paracingulate sulcus than healthy controls and also to non-hallucinating patients, and suggested that this region of the brain is tuned to “reality monitoring”, i.e. the ability to judge whether a memory comes from an outer or inner source. This suggestion^22^ was based on the findings reported by Buda, et al.^23^ that otherwise healthy individuals, born without a discernible paracingulate sulcus in either hemisphere, showed impaired performance on a word-pair memory/imagery task. These observations^22, 23^ may provide important clues for understanding the significance of the present findings, insofar that the ventral portion of this cortical region may be crucial for how AVHs are spontaneously initiated and also why they may spontaneously and transiently disappear. In this respect our data support a previous report from two patients of anticipatory neural activity to onset-signaling^24^. A potential confounding of the results could be anticipatory attention focus on motor-responses (cf.^25^), which otherwise could affect the observed activity. As seen in the lower panels of Figure 3 this is probably not the case, since there was a clear peak around 5 seconds post-response obtained from the pre-central motor-cortex on the left side (from right-handheld response buttons), and with 5-6 times higher response amplitude as that obtained from the ventro medial frontal cortex^26^. This is what one would expect considering the lag of the hemodynamic response relative to a neural event, but would not expect for a peak occurring at 3-4 seconds and in the ventro-medial frontal cortex, after the button-press. The decrease in brain activity a few seconds before the indicated awareness of the offset of a hallucinatory episode, may correspond to previous findings of frontal neurotransmitter imbalance^27–29^. Using MR spectroscopy (^1^H-MRS), Ćurčić-Blake, et al.^30^, van Den Heuvel, et al.^31^ and Hugdahl, et al.^32^ found increased levels of glutamate in frontal regions in hallucinating individuals. This is in accordance with what Jardri, et al.^28^ labelled the Excitatory/Inhibitory (E/I) imbalance model of auditory hallucinations, and we now suggest that the offset of a hallucinatory event is mediated by temporary restoring such imbalances. Future research will hopefully resolve the underlying causes at the receptor and transmitter level. The present findings could also have therapeutic implications in guiding more targeted brain stimulation approaches. Brain stimulation interventions is a promising approach to medication-resistant hallucinations^33,34,35^ and targeting the the ventro-medial frontal cortex using stimulation to stop the onset, or accelerate the offset of an AVH-episode could be a way to help patients overcome intractable AVHs.

## Acknowledgements

The authors want to acknowledge the contribution of MR-technicians and research assistants, as well as the clinical and non-clinical participants, at each site.

## Methods

### Subjects

Structural data and functional BOLD data were collected from a total of 66 subjects, of which 45 were diagnosed with an ICD-10 or DSM-IV schizophrenia spectrum disorder. The patients came from three collaborating projects and sites. These were University of Bergen, Norway (n=11, 7 males, mean age 27.8 (SD 7.0) years); Plovdiv Medical University, Bulgaria (n=13, 11 males, mean age 35.3 (SD 14.0 years), Groningen University Medical Center, Netherlands (n=21, 7 males, mean age 39.0 (SD 11.4) years), total 25 males and 19 females, mean age 37.9 (SD 13.2) years. Symptom severity for patients was assessed with the positive and negative syndrome scale (PANSS^36^). Mean total PANSS score was 64.9 (SD 16.9). In order to be included in the study, patients had to score >3 on the PANSS, P3 hallucinatory behavior item within a week of the MR scanning (mean P3 score 4.6 (SD 1.1)). The patients were all on second-generation antipsychotics (often clozapine), with some patients in addition being prescribed antidepressants and/or anxiolytics. Mean antipsychotic Defined Daily Doses (DDD) were 1.228 (SD 0.578). The total sample also consisted of 21 non-clinical hallucinating individuals (4 males, mean age 44.5 (SD 13.0) years), i.e. in whom a clinical axis I and axis II diagnosis was ruled out using the CASH and SCID-II interview, included in the Groningen University Medical Center sample (for details see^10^). This yielded a total sample of 29 males, 37 females, mean age 38.2 (SD 13.0) years. The study was approved by the local ethics committees at each site, and had a European Research Council Ethics Approval (ERCEA 2016-439428). Transfer of data between the sites and re-analysis at the Bergen University was approved by the ethical committees at each site, and further confirmed by the Regional Committee for Medical Research Ethics in Western Norway (REK-Vest #2017/933).

### Data collection

Functional MR data were collected with a “symptom-capture” paradigm^9^, where subjects were instructed to press a button when a hallucinatory episode began (onset), and to press another button when the episode ended (offset). The instruction of when to press the buttons was presented visually through LCD goggles mounted on the head-coil, in the language appropriate to the location, along with a fixation cross that was displayed in the middle of the visual field. A time-window was set from 10 seconds before to 15 seconds after the subject had pressed a button, from which voxel-wise data were extracted, analyzed and displayed in a sliding window over the evaluation period. Particulars of the acquisition varied between sites. Gradient Echo Planar Imaging (EPI) was used to collect functional BOLD data on all sites. Data from Bergen were collected using a 3T GE 750 scanner with a 32-, or 8-channel head coil (300 volumes at TR = 2000 ms, total duration 10 min, TE = 30 ms, flip angle 90°, resolution 128 × 128, pixel spacing 1.72 mm, 30 or 26 slices of 3mm thickness with 0.5mm gap). Plovdiv data were collected with a 3T GE 750w scanner and 24-channel head coil (900 volumes at TR=2000 ms (total duration 30 min), TE = 30 ms, flip angle 90°, resolution 64 × 64, pixel spacing 3.44 mm, 34 slices of 3 mm thickness with 0.5 mm gap). Groningen data were acquired on a 3T Philips Achieva scanner, as 800 volumes at TR = 21.75 ms (total duration 8 min, 6 sec), TE = 32.4 ms, 64 × 64, 4 mm voxel size, 40 slices (4 mm thickness), no gap. This scan sequence achieves full brain coverage within 609 ms by combining a 3D-PRESTO pulse sequence with parallel imaging (SENSE) in two directions using a commercial 8-channel SENSE head-coil. A high-resolution structural T1 volume was acquired for each subject, along with additional sequences that varied between sites, of no relevance and are not reported herein.

### Motor (button-press) response data

Upon visual inspection of the subjects’ button-press data, it was found that a certain number of subjects reported multiple episode-onsets which did not distinctly match to a single end-of- “voice” event, which required interpretation and operational definition of episodes. We interpreted and operationally defined the relationship between onset- and offset button responses in three different ways, described and illustrated in Figure 5 below. According to each definition, subjects’ button-press response-data were filtered to remove spurious episodes, and extract distinct blocks of hallucinatory vs non-hallucinatory periods. Additional criteria regarding minimal spacing between events ensured the validity of the subsequent analyses.

**Figure 4.**
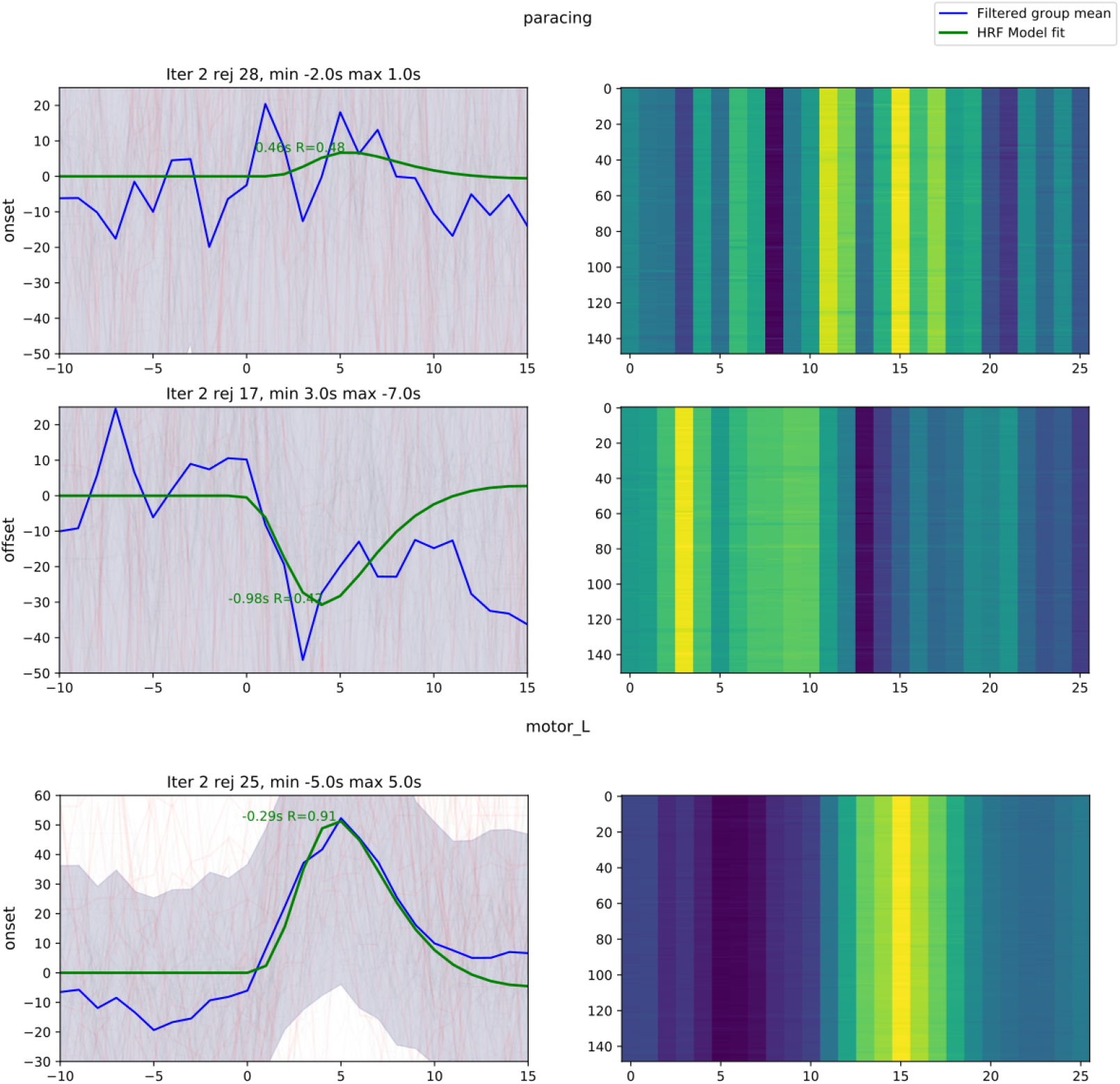
shows group-average (blue line) and hemodynamic response function (HRF) - model fit (green line) of time-course data from the nominated ROI in the intersection of the paracingulate cortex/medial inferior frontal cortex and the frontal pole, separated for onset (left upper panel) and offset (left middle panel) events, for the final iteration of filtering. The lower panel show corresponding time-courses for the left pre-central motor cortex (note different y-axis scale). Adjacent panels to the right show group-mean encoded as intensity, for each step (y-axis) of a leave-one-out validation according to which strongly deviant time courses were rejected. See Methods and Results sections for further details.

**Figure 5.**
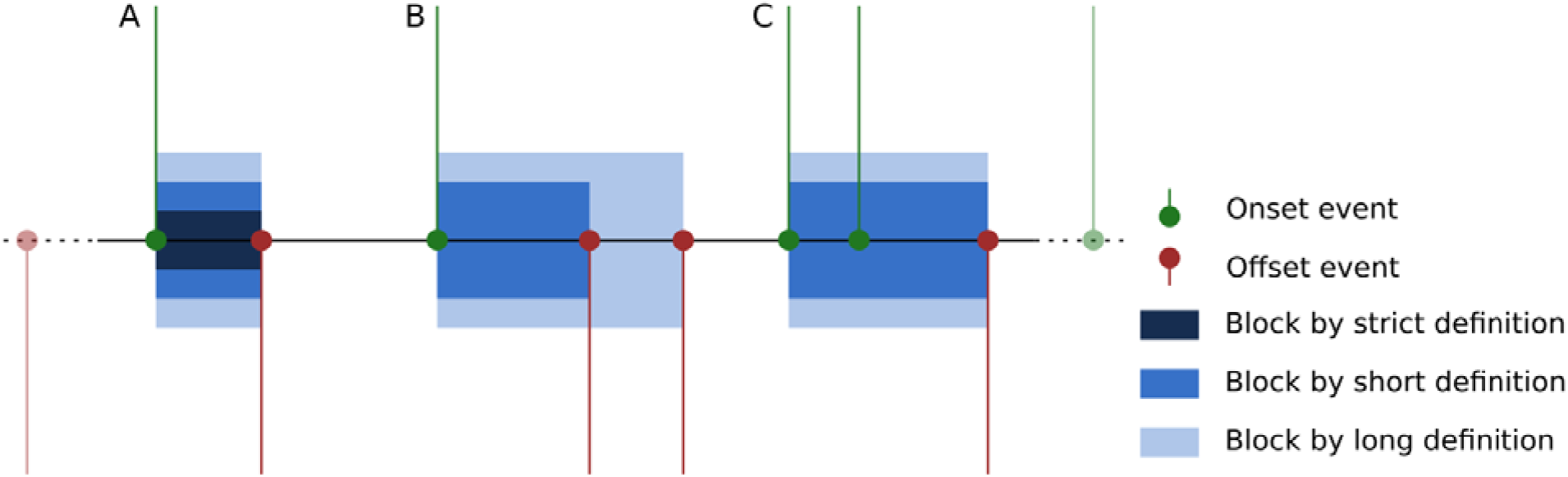
illustrates the three possible interpretations of subjects’ response patterns. Pattern A is unambiguous, and is matched identically by strict, short and long definitions. Pattern B includes a spurious “offset” event; the strict definition rejects this block, the short definition ends the block at the first offset, and the long definition ends the block at the last offset. Pattern C shows spurious onset events; again this block was rejected by the strict definition, whilst both short and long definitions match from the first “onset” event.

#### Operational definition of hallucinatory and non-hallucinatory periods

According to a first possible interpretation and definition, denoted “short blocks”, hallucinatory periods were defined as spanning from the moment a subject first pressed the button indicating the start-of-voice episode, until they first pressed the button to indicate the end-of-voice episode. This yielded 1055 usable blocks across all subjects. In a second interpretation, denoted “long blocks”, hallucinatory periods were defined as spanning from the moment a subject first pressed the button indicating the start-of-voice episode, until the last press indicating the end-of-voice episode before the next reported start-of-voice episode. This yielded 1000 usable blocks across all subjects. The final, “strict blocks” interpretation, accepted only periods unambiguously bounded by a single start-of-voice and a single end-of-voice button-press. This yielded 300 usable blocks across all subjects. Figure 5 shows the different interpretations, illustrated on a representative subject’s response data.

### Pre-processing of fMRI data

Functional MR data were pre-processed using standard tools from FSL 5.0.11 (FEAT pipeline), with additional filtering for artifacts using the ICA-AROMA method^37–40^. Brain masks for each subject’s structural T1-images and functional data were derived using FSL’s bet2 tool, with fractional intensity threshold and initial mesh center-of-gravity tuned on a group basis to accommodate differences between the three sites. Individual brain-masks were manually subject to visual quality control to ensure completeness and specificity of the mask. When necessary, brain masking was repeated for a small number of subjects using individually tuned parameters. Next, individual brain-extracted functional data were subject to motion correction, using FSL’s MCFLIRT utility^38, 41^. Data were filtered spatially with a 5 mm FWHM Gaussian kernel, and temporally with a 100 sec high-pass filter. They were then registered to individual, brain-extracted high-resolution structural images using FSL’s linear registration tool (FLIRT), with the recommended Boundary-Based Registration mode and a restricted (35 degree) search range. Registration was inspected visually for validity; for three subjects where the BBR method performed poorly, a 12-degree-of-freedom registration was substituted. Individual brain-extracted high-resolution structural images were registered to a standard 2mm T1 template in MNI152 space (ICBM152, non-linear, 6^th^ generation), with a 12-degrees-of-freedom linear registration using FSL FLIRT, followed by a nonlinear registration with 10 mm warp resolution using FSL FNIRT. The ICA-AROMA method was applied to filtered, native-space functional data to remove residual signals associated with motion artifacts and noise. Finally, per-subject native-space functional masks were transformed into standard space using non-linear parameters derived from the registration steps; these masks were assessed programmatically to ensure adequate coverage of the brain, with particular attention to prescribed regions of interest (ROIs), see below.

### Block analysis

To confirm the validity of the filtered response data, a standard fMRI block--analysis was performed using FSL FEAT first-level and higher-level analysis pipelines^42^, applied for the long block definition. Mixed effects modeling (FLAME 1+2) was used for higher-level analysis; clusters were thresholded non-parametrically at Z>4.5; corrected cluster significance thresholded at p>0.05 and extent > 20 voxels.

### Time-course analysis

For each hallucinatory period, identified as demarcating the start or end of a hallucinatory episode, windows of functional data were extracted extending from a time of t = −10 sec through to t = +15 sec, relative to the moment in time at which the button-press event occurred (set to t = 0 sec), as a continuous variable. Since the temporal resolution of functional data is relatively coarse, and also varied between sites in the current dataset, data were sampled at regular 1sec intervals, weighted and normalized according to a Gaussian kernel in the temporal dimension (FWHM = 0.94 sec). An initial principal component analysis (PCA) on grouped, extracted functional segments, guided the selection of ROIs for further inspection and analysis. Cluster locations identified in the functional analysis and those nominated and reported in previous meta-analyses^16, 17^ were also inspected for overlaps of activity between the current study and activity reported in the meta-analyses (see Figure 1 and Table 1). For each nominated ROI, and for each of the extracted functional segments, time-course vectors were obtained and spatially averaged over a 5 mm radius sphere, allowing activity in each region to be examined and evaluated in the time-frame leading up to and immediately following a button-press, marking the onset and offset of a hallucinatory episode. Separately for onset and offset events, time-course vectors for each region were aligned in the temporal dimension to the group-average for the respective region (maximizing cross-correlation). Aberrant time-courses (correlation varying by more than two standard deviations) were rejected, iteratively updating the group to refine an estimated model time course for the region around onset and offset events as shown in Figure 4. A dual-gamma hemodynamic response-function (HRF) model was thereafter fitted to the refined time-course model, allowing magnitude and timing of any activity-related peak to be identified and statistically evaluated. Random permutation-analysis (n = 5000) was performed to identify differential effects on fit parameters between onset- and offset-events.

### 4D permutation analysis

The ROI-based analysis was generalized to a full four-dimensional (4D) permutation analysis, characterizing activity specific to onset- or offset-events at each time-point in the extracted window segments, searching across the entire brain volume voxel-wise. Due to the large volume of data and computationally intensive nature of the permutation analysis, it was necessary to develop a new software tool to facilitate this analysis. P-values were extracted (n = 10,000 permutations), along with p-values calculated on a gamma approximation of the obtained distribution^43^, for each voxel, at each time-point. Initially, time-windows associated with onset- and offset-events were contrasted jointly in the permutation analysis, yielding differential effects for onset and offset events. Subsequently, time-points from offset- and onset-events were separately contrasted against windows extracted around random time points (without synchronization to subjects’ button-responses), as a baseline state.

## Data availability statement

The datasets analysed during the current study are not publicly available due to restrictions imposed by Regional Committee for Medical Research Ethics in Western Norway (REK Vest) and the Data Protection Officer of the Western Norway Health Authorities (Personvernombudet) but are available from the corresponding author on reasonable request

## Software/Code availability statement

Novel in-house developed software implemented for this study has been made publicly available here: https://git.app.uib.no/bergen-fmri/functional-transients. All other stages of analysis were performed using widely-available open-source software, including tools from the FSL suite and additional filtering with the ICA-AROMA method.

## Author contributions

KH designed the study, participated in patient recruitment, data analysis and interpretation, wrote the ms, ARC analyzed the data, participated in data acquisition and interpretation, commented on the ms, EJ participated in patient recruitment, commented on the ms, LE participated in data acquisition and analysis, commented on the ms, DS participated in patient recruitment. commented on the ms, SK participated in patient recruitment, commented on the ms, LBS participated in organization of data and ms and commented on the ms, RAK participated in patient recruitment, commented on the ms, EML participated in patient recruitment, commented on the ms, IES participated in patient and subjects recruitment, data interpretation, and commented on the ms.

## Supplementary information

Correspondence and requests for materials should be addressed to Kenneth Hugdahl.

## Acknowledgments

The authors want to acknowledge the contribution by MR-technicians and participants for making the study possible.

## References

1 Andreasen, N. C. & Olsen, S. Negative v positive schizophrenia. Definition and validation. JAMA Psychiatry 39, 789–794, doi:10.1001/archpsyc.1982.04290070025006 (1982).

2 Waters, F., Badcock, J., Michie, P. & Maybery, M. Auditory hallucinations in schizophrenia: Intrusive thoughts and forgotten memories. Cogn. Neuropsychiatry 11, 65–83, doi:10.1080/13546800444000191 (2006).

3 Aleman, A. & Larøi, F. Hallucinations: The science of idiosyncratic perception. (American Psychological Association, 2008).

4 Sartorius, N., Jablensky, A., Korten, A., Ernberg, G., Anker, M., Cooper, J. E. & Day, R. Early manifestations and first-contact incidence of schizophrenia in different cultures. A preliminary report on the initial evaluation phase of the WHO collaborative study on determinants of outcome of severe mental disorders. Psychol. Med. 16, 909–928, doi:10.1017/s0033291700011910 (1986).

5 Ford, J. M., Morris, S. E., Hoffman, R. E., Sommer, I. E., Waters, F., McCarthy-Jones, S., Thoma, R. J., Turner, J. A., Keedy, S. K., Badcock, J. C. & Cuthbert, B. N. Studying Hallucinations within the NIMH RDoC Framework. Schizophr. Bull. 40, 295–304, doi:10.1093/schbul/sbu011 (2014).

6 Ford, J. M., Dierks, T., Fisher, D. J., Herrmann, C. S., Hubl, D., Kindler, J., Koenig, T., Mathalon, D. H., Spencer, K. M., Strik, W. & van Lutterveld, R. Neurophysiological studies of auditory verbal hallucinations. Schizophr. Bull. 38, 715–723, doi:10.1093/schbul/sbs009 (2012).

7 Hugdahl, K. & Sommer, I. E. Auditory verbal hallucinations in schizophrenia from a levels of explanation perspective. Schizophr. Bull. 44, 234–241, doi:10.1093/schbul/sbx142 (2018).

8 Kråkvik, B., Larøi, F., Kalhovde, A. M., Hugdahl, K., Kompus, K., Salvesen, Ø., Stiles, T. C. & Vedul□Kjelsås, E. Prevalence of auditory verbal hallucinations in a general population: A group comparison study. Scand. J. Psychol. 56, 508–515, doi:10.1111/sjop.12236 (2015).

9 Sommer, I. E., Diederen, K. M., Blom, J.-D., Willems, A., Kushan, L., Slotema, K., Boks, M. P. M., Daalman, K., Hoek, H. W., Neggers, S. F. W. & Kahn, R. S. Auditory verbal hallucinations predominantly activate the right inferior frontal area. Brain 131, 3169–3177, doi:10.1093/brain/awn251 (2008).

10 Sommer, I. E., Daalman, K., Rietkerk, T., Diederen, K. M., Bakker, S., Wijkstra, J. & Boks, M. P. M. Healthy individuals with auditory verbal hallucinations; Who are they? Psychiatric assessments of a selected sample of 103 subjects. Schizophr. Bull. 36, 633–641, doi:10.1093/schbul/sbn130 (2010).

11 Larøi, F., Sommer, I. E., Blom, J. D., Fernyhough, C., ffytche, D. H., Hugdahl, K., Johns, L. C., McCarthy-Jones, S., Preti, A., Raballo, A., Slotema, C. W., Stephane, M. & Waters, F. The characteristic features of auditory verbal hallucinations in clinical and nonclinical groups: State-of-the-art overview and future directions. Schizophr. Bull. 38, 724–733, doi:10.1093/schbul/sbs061 (2012).

12 Daalman, K., Boks, M. P. M., Diederen, K. M., de Weijer, A. D., Blom, J. D., Kahn, R. S. & Sommer, I. E. The same or different? A phenomenological comparison of auditory verbal hallucinations in healthy and psychotic individuals. J. Clin. Psychiat 72, 320–325, doi:10.4088/jcp.09m05797yel (2011).

13 Bless, J. J., Larøi, F., Kompus, K., Kråkvik, B., Vedul-Kjelsås, E., Kalhovde, A. M. & Hugdahl, K. Do adverse life events at first onset of auditory verbal hallucinations influence subsequent voice characteristics? Results from an epidemiological study. Psychiatry Res. 261, 232–236, doi:10.1016/j.psychres.2017.12.060 (2017).

14 Delespaul, P., deVries, M. & van Os, J. Determinants of occurrence and recovery from hallucinations in daily life. Soc. Psychiatry Psychiatr. Epidemiol. 37, 97–104, doi:10.1007/s001270200000 (2002).

15 Moonen, C. T. W. & Bandettini, P. A. Functional MRI. (Springer-Verlag, 1999).

16 Jardri, R., Pouchet, A., Pins, D. & Thomas, P. Cortical activations during auditory verbal hallucinations in schizophrenia: A coordinate-based meta-analysis. Am. J. Psychiatry 168, 73–81, doi:10.1176/appi.ajp.2010.09101522 (2011).

17 Kompus, K., Westerhausen, R. & Hugdahl, K. The “paradoxical” engagement of the primary auditory cortex in patients with auditory verbal hallucinations: A metaanalysis of functional neuroimaging studies. Neuropsychologia 49, 3361–3369, doi:10.1016/j.neuropsychologia.2011.08.010 (2011).

18 Dierks, T., Linden, D. E., Jandl, M., Formisano, E., Goebel, R., Lanfermann, H. & Singer, W. Activation of Heschl’s gyrus during auditory hallucinations. Neuron 22, 615–621, doi:10.1016/s0896-6273(00)80715-1 (1999).

19 van de Ven, V. G., Formisano, E., Röder, C. H., Prvulovic, D., Bittner, R. A., Dietz, M. G., Hubl, D., Dierks, T., Federspiel, A., Esposito, F., Di Salle, F., Jansma, B., Goebel, R. & Linden, D. E. J. The spatiotemporal pattern of auditory cortical responses during verbal hallucinations. Neuroimage 27, 644–655, doi:10.1016/j.neuroimage.2005.04.041 (2005).

20 Eichele, T., Debener, S., Calhoun, V., Specht, K., Engel, A., Hugdahl, K., Von Cramon, D. Y. & Ullsperger, M. Prediction of human errors by maladaptive changes in event-related brain networks. Proc. Natl. Acad. Sci. U. S. A. 105, 6173–6178, doi:10.1073/pnas.0708965105 (2008).

21 Alonso-Solís, A., Vives-Gilabert, Y., Grasa, E., Portella, M. J., Rabella, M., Sauras, R. B., Roldán, A., Núñez-Marín, F., Gómez-Ansón, B., Pérez, V., Alvarez, E. & Corripio, I. Resting-state functional connectivity alterations in the default network of schizophrenia patients with persistent auditory verbal hallucinations. Schizophr. Res. 161, 261–268, doi:10.1016/j.schres.2014.10.047 (2015).

22 Garrison, J. R., Fernyhough, C., McCarthy-Jones, S., Simons, J. S. & Sommer, I. E. Paracingulate sulcus morphology and hallucinations in clinical and nonclinical groups. Schizophr. Bull. 45, 733–741, doi:10.1093/schbul/sby157 (2019).

23 Buda, M., Fornito, A., Bergström, Z. M. & Simons, J. S. A specific brain structural basis for individual differences in reality monitoring. J. Neurosci. 31, 14308–14313, doi:10.1523/jneurosci.3595-11.2011 (2011).

24 Shergill, S. S., Brammer, M., Amaro, E., Williams, S., Murray, R. & Mcguire, P. Temporal course of auditory hallucinations. Br. J. Psychiatry 185, 516–517, doi:10.1192/bjp.185.6.516 (2004).

25 van Lutterveld, R., Diederen, K., Schutte, M., Bakker, R., Zandbelt, B. & Sommer, I. E. Brain correlates of auditory hallucinations: Stimulus detection is a potential confounder. Schizophr. Res. 150, 319–320, doi:10.1016/j.schres.2013.07.021 (2013).

26 Logothetis, N. K. & Pfeuffer, J. On the nature of the BOLD fMRI contrast mechanism. Magn. Reson. Imaging 22, 1517–1531, doi:10.1016/j.mri.2004.10.018 (2004).

27 Steinmann, S., Leicht, G. & Mulert, C. The interhemispheric miscommunication theory of auditory verbal hallucinations in schizophrenia. Int. J. Psychophysiol. 145, 83–90, doi:10.1016/j.ijpsycho.2019.02.002 (2019).

28 Jardri, R., Hugdahl, K., Hughes, M., Brunelin, J., Waters, F., Alderson-Day, B., Smailes, D., Sterzer, P., Corlett, P. R., Leptourgos, P., Debbané, M., Cachia, A. & Denève, S. Are hallucinations due to an imbalance between excitatory and inhibitory influences on the brain? Schizophr. Bull. 42, 1124–1134, doi:10.1093/schbul/sbw075 (2016).

29 Allen, P., Sommer, I. E., Jardri, R., Eysenck, M. W. & Hugdahl, K. Extrinsic and default mode networks in psychiatric conditions: Relationship to excitatory-inhibitory transmitter balance and early trauma. Neurosci. Biobehav. Rev. 99, 90–100, doi:10.1016/j.neubiorev.2019.02.004 (2019).

30 Ćurčić-Blake, B., Bais, L., Sibeijn-Kuiper, A., Pijnenborg, H. M., Knegtering, H., Liemburg, E. & Aleman, A. Glutamate in dorsolateral prefrontal cortex and auditory verbal hallucinations in patients with schizophrenia: A 1H MRS study. Prog. Neuro-Psychopha 78, 132–139, doi:10.1016/j.pnpbp.2017.05.020 (2017).

31 van Den Heuvel, M. P., Mandl, R. C. W., Stam, C. J., Kahn, R. S. & Hulshoff Pol, H. E. Aberrant frontal and temporal complex network structure in schizophrenia: A graph theoretical analysis. J. Neurosci. 30, 15915–15926, doi:10.1523/jneurosci.2874-10.2010 (2010).

32 Hugdahl, K., Craven, A. R., Nygård, M., Løberg, E.-M., Berle, J.Ø., Johnsen, E., Kroken, R., Specht, K., Andreassen, O. A. & Ersland, L. Glutamate as a mediating transmitter for auditory hallucinations in schizophrenia: A 1H MRS study. Schizophr. Res. 161, 252–260, doi:10.1016/j.schres.2014.11.015 (2015).

33 Stoyanov, D., Stieglitz, R.-D., Lenz, C. & Borgwardt, S. in New developments in clinical psychology research (eds D. Stoyanov & R-D. Stieglitz) 195–209 (Nova Science Publishers, New York, 2015).

34 Dougall, N., Maayan, N., Soares-Weiser, K., McDermott, L. M. & McIntosh, A. Transcranial magnetic stimulation (TMS) for schizophrenia. Cochrane Database Syst. Rev., doi:10.1002/14651858.cd006081.pub2 (2015).

35 Nathou, C., Simon, G., Dollfus, S. & Etard, O. Cortical anatomical variations and efficacy of rTMS in the treatment of auditory hallucinations. Brain Stimul. 8, 1162–1167, doi:10.1016/j.brs.2015.06.002 (2015).

## Methods references

36 Kay, S. R., Fiszbein, A. & Opler, L. A. The positive and negative syndrome scale (PANSS) for schizophrenia. Schizophr. Bull. 13, 261–276, doi:10.1093/schbul/13.2.261 (1987).

37 Pruim, R. H. R., Mennes, M., van Rooij, D., Llera, A., Buitelaar, J. K. & Beckmann, C. F. ICA-AROMA: A robust ICA-based strategy for removing motion artifacts from fMRI data. Neuroimage 112, 267–277, doi:10.1016/j.neuroimage.2015.02.064 (2015).

38 Jenkinson, M. & Smith, S. A global optimisation method for robust affine registration of brain images. Med. Image Anal. 5, 143–156, doi:10.1016/S1361-8415(01)00036-6 (2001).

39 Woolrich, M. W., Ripley, B. D., Brady, M. & Smith, S. M. Temporal autocorrelation in univariate linear modeling of fMRI data. Neuroimage 14, 1370–1386, doi:10.1006/nimg.2001.0931 (2001).

40 Jenkinson, M., Beckmann, C. F., Behrens, T. E. J., Woolrich, M. W. & Smith, S. M. FSL. Neuroimage 62, 782–790, doi:10.1016/j.neuroimage.2011.09.015 (2012).

41 Jenkinson, M., Bannister, P., Brady, M. & Smith, S. Improved optimization for the robust and accurate linear registration and motion correction of brain images. Neuroimage 17, 825–841, doi:10.1006/nimg.2002.1132 (2002).

42 Woolrich, M. W., Behrens, T. E. J., Beckmann, C. F., Jenkinson, M. & Smith, S. M. Multilevel linear modelling for fMRI group analysis using Bayesian inference. Neuroimage 21, 1732–1747, doi:10.1016/j.neuroimage.2003.12.023 (2004).

43 Winkler, A. M., Ridgway, G. R., Douaud, G., Nichols, T. E. & Smith, S. M. Faster permutation inference in brain imaging. Neuroimage 141, 502–516, doi:10.1016/j.neuroimage.2016.05.068 (2016).

